# Phosphosite T674A mutation in kinesin family member 3A fails to reproduce tissue and ciliary defects characteristic of CILK1 loss of function

**DOI:** 10.1101/2020.06.30.180307

**Authors:** Casey D. Gailey, Eric J. Wang, Li Jin, Sean Ahmadi, David L. Brautigan, Xudong Li, Wenhao Xu, Zheng Fu

**Author notes:** Correspondence (Z.F.); (Z.F.). Funding information: National Institute of General Medical Sciences, Grant Number: GM127690 (to ZF); National Institute of arthritis and musculoskeletal and skin diseases, Grant Number: AR064792 (to XL).

## Abstract

**Background:** Kinesin family member 3A (KIF3A) is a molecular motor protein in the heterotrimeric kinesin-2 complex that drives anterograde intraflagellar transport. This process plays a pivotal role in both biogenesis and maintenance of the primary cilium that supports tissue development. Ciliogenesis associated kinase 1 (CILK1) phosphorylates human KIF3A at Thr672. CILK1 loss of function causes ciliopathies that manifest profound and multiplex developmental defects, including polydactyly, shortened and hypoplastic bones and alveoli airspace deficiency, leading to perinatal lethality. Prior studies have raised the hypothesis that CILK1 phosphorylation of KIF3A is critical for its regulation of organ development.

**Results:** We produced a mouse model with phosphorylation site Thr674 in mouse *Kif3a* mutated to Ala. *Kif3a* T674A homozygotes are viable and exhibit no skeletal abnormalities, and only mildly reduced airspace in alveoli. Mouse embryonic fibroblasts carrying *Kif3a* T674A mutation show a normal rate of ciliation and a moderate increase in cilia length.

**Conclusion:** These results indicate that eliminating Kif3a Thr674 phosphorylation by CILK1 is insufficient to reproduce the severe developmental defects in ciliopathies caused by Cilk1 loss of function. This suggests KIF3A phosphorylation by CILK1 is not essential for tissue development and other substrates are involved in Cilk1 ciliopathies.

## 1. INTRODUCTION

Most eukaryotic cells use the primary cilium as the sensory organelle and signaling antenna to detect environmental cues and transmit extracellular signals to modulate intracellular processes and cell behaviors.^1^ Primary cilia are enriched with membrane receptors, ion channels, metabolic enzymes, and second messengers, and provide a highly efficient and intricate infrastructure for the operation of many signaling pathways such as Hedgehog, G protein-coupled receptors (GPCR), and receptor tyrosine kinases (RTK).^2,3^ A variety of human genetic disorders, collectively called ciliopathies, is caused by mutations in ciliary proteins that disrupt the integrity of cilia structure and signaling.^4^ A deeper comprehension of the molecular mechanism regulating cilia structure and function will be instrumental for a better understanding of the pathogenesis of ciliopathies.

Ciliogenesis associated kinase 1 (CILK1),^5^ formerly known as intestinal cell kinase (ICK),^6^ is emerging as a key protein localized at the cilia base that restricts cilia biogenesis and length.^7-13^ Human mutations in *CILK1* have been linked to ciliopathies and epilepsy.^10,14-16^ Both *Cilk1* null and mutated (R272Q) knock-in mouse models have shown that CILK1 dysfunction causes perinatal lethality and detrimental effects on multi-organ development, resulting in anomalies such as hydrocephalus, alveoli airspace deficiency, polydactyly, shortened and hypoplastic bones.^7,8,10,11,17^ Although the essential role for CILK1 in development and primary cilia has been established, we still do not know much about which CILK1 substrates mediate its actions on developmental and ciliary phenotype. Among several candidate substrates identified for CILK1, kinesin family member 3A (KIF3A) has gained significant attention because its critical role as a motor protein in ciliary transport is directly related to the control of cilia formation.^7,12^

Intraflagellar transport (IFT) along cilia axonemal microtubules consists of kinesin-mediated anterograde and dynein-mediated retrograde transport. Kinesin 3A/3B dimeric motor subunits and kinesin-associated protein (KAP) form the heterotrimeric Kinesin-2 complex.^18^ Anterograde IFT mediated by KIF3A/KIF3B/KAP is essential for the assembly and maintenance of primary cilia.^19,20^ The C-terminal cargo-binding domain of human KIF3A contains multiple phosphorylation sites, including PKA site Ser687, CaMKII sites Thr692 and Ser696, and CILK1 site Thr672. Phosphorylation of S687/T692/S696 enhances the cargo-binding and trafficking activities of KIF3A.^21^ The CILK1 site Thr672 in human KIF3A is highly conserved among metazoans. CILK1 robustly phosphorylates human KIF3A-Thr672 (mouse Kif3a-Thr674) in vitro and in vivo.^7,12^ A previous study indicates that mouse Kif3a-T674A mutant protein has a stronger capability than Kif3a-WT protein to rescue cilia formation in Kif3a-knockdown cells, suggesting a restrictive effect of Kif3a-Thr674 phosphorylation on cilia assembly.^7^ Based on these pieces of evidence, we postulated that phosphorylation of KIF3A-Thr672 acts as an important downstream effector through which CILK1 restricts cilia assembly and regulates organ development. To test this hypothesis, we generated a *Kif3a*-T674A knock-in mouse model and interrogated the developmental and ciliary phenotype caused by phospho-Thr674 deficiency in Kif3a.

## 2. RESULTS

### 2.1 Developmental phenotype of *Kif3a* T674A knock-in mice

We used CRISPR/Cas9 to genetically engineer several mutations in exon 17 of mouse *Kif3a* (Fig. 1A). These knock-in mutations enable not only the conversion of Thr674 to Ala but also the introduction of a StuI restriction enzyme recognition site near the T674A mutation, as confirmed by sequencing results (Fig. 1B). Restriction digestion of PCR products by StuI allowed us to distinguish between *Kif3a* wild type (WT) and mutant alleles, as shown in genotyping results (Fig. 1C). We then evaluated whether this mutation affects Kif3a protein stability and subcellular localization in mouse embryonic fibroblasts. Similar levels of Kif3a WT and T674A mutant proteins were detected in whole cell extracts, suggesting the T674A mutation did not alter KIF3A stability (Fig. 1B). Kif3a WT protein was enriched in the perinuclear region, showing punctate staining in a reticular pattern. A fraction of Kif3a resided at the base of the primary cilium (Fig. 1C). Kif3a T674A mutant protein did not appear to be mislocalized; there was not a distinct difference between Kif3a mutant and WT proteins in their subcellular distribution (Fig. 1D-E).

**Figure 1:**
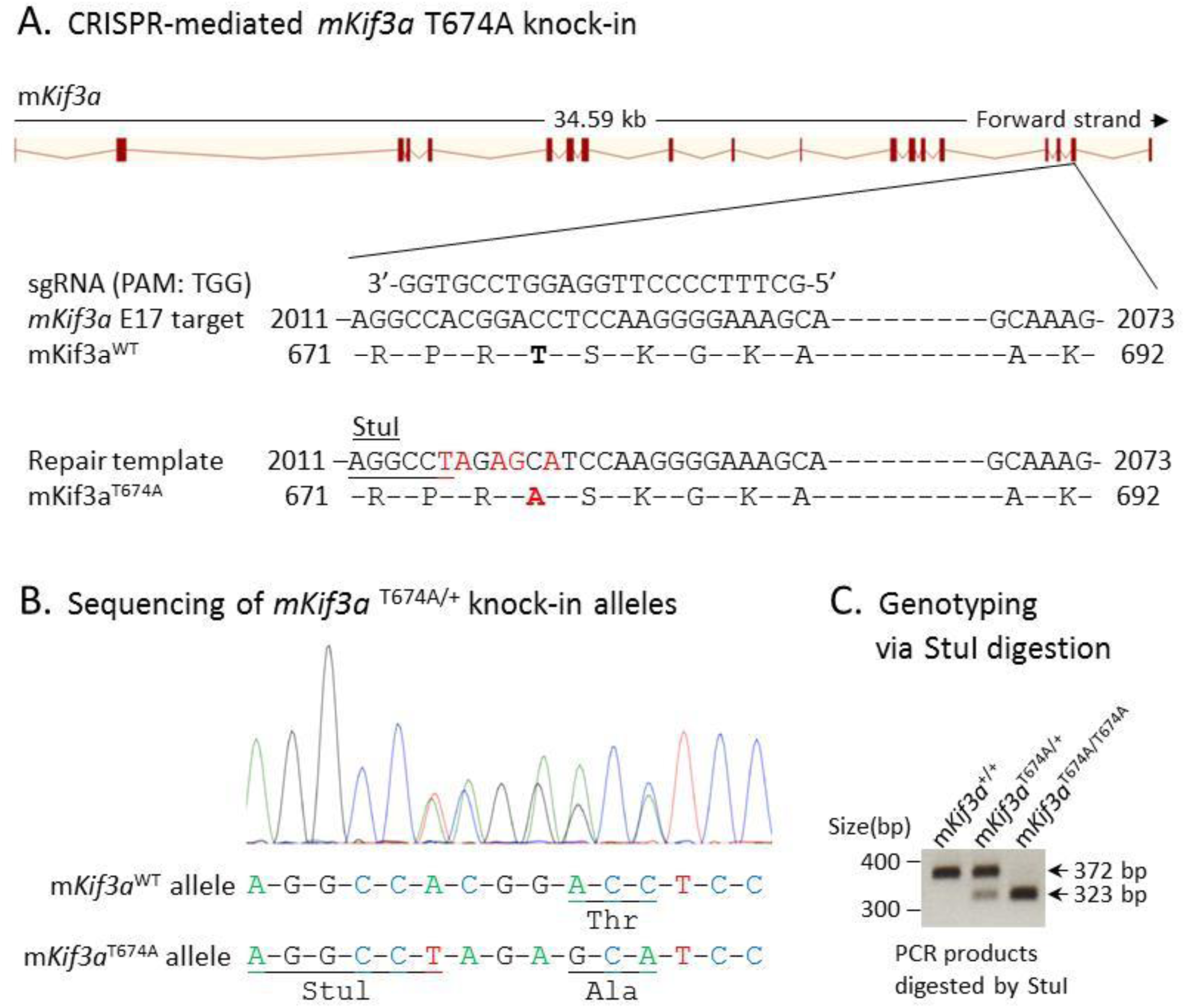
Generation of the *Kif3a* T674A knock-in mouse model. (A) A schematic illustration of CRISPR/Cas9-mediated knock-in of the phospho-deficient mutation T674A in *mKif3a*. A StuI restriction enzyme cutting site was also introduced into the *mKif3a*^T674A^ mutant allele by a silent mutation (CCA→CCT). (B) Sequencing results confirming designed mutations that not only convert Thr674 to Ala but also introduce a new StuI site near T674A for genotyping. Two silent base substitutions (CGG→AGA) were also engineered to deter the recut of the repair template by Cas9 nuclease. (C) Genotyping results from StuI-digested PCR products that can distinguish between *mKif3a*^+/+^ (wild type), *mKif3a*^T674A/+^ (heterozygotes) and *mKif3a*^T674A/T674A^ (homozygotes) littermates.

Mice bearing *Cilk1* homozygous null or kinase deficient (R272Q) mutations died at birth and showed profound developmental abnormalities in multi-organ systems.^7,8,10,11,17^ The perinatal lethality phenotype of these Cilk1 mutant mice was linked to severe alveolar airspace deficiency in the lung.^11^ In contrast, mice bearing *Kif3a* T674A mutations were viable. Compared with wild type lungs, Kif3a homozygous, but not heterozygous, mutant lungs showed only a moderate decrease in alveolar airspace (Fig. 2). Cilk1 homozygous mutant mice also displayed extensive skeletal defects, including polydactyly, shortened limbs, and deformed spine.^10,17^ However, these aberrations were not observed in whole-mount skeletal staining (Fig. 3) and full-body X-Ray scan (data not shown) of Kif3a T674A mutant mice. We concluded that absence of Kif3a phosphorylation at Thr674 produced only mild airspace deficiency in the lung and was insufficient to reproduce the profound anomalies in the skeleton caused by CILK1 loss of function.

**Figure 2:**
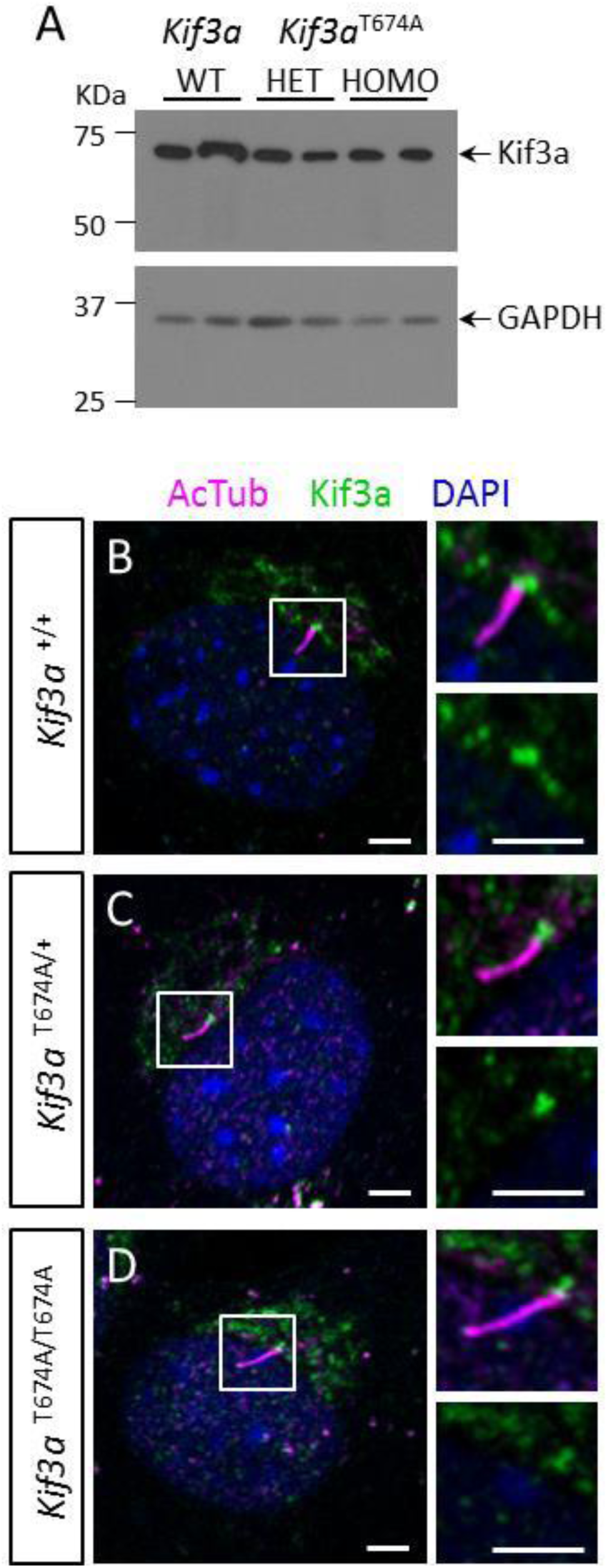
Kif3a protein expression and subcellular localization in Kif3a wild type and T674A mutant mouse embryonic fibroblasts. (A) Western blots show Kif3a and GAPDH (loading control) expression in whole cell extracts of Kif3a wild type (WT), T674 heterozygous (HET) and homozygous (HOMO) mouse embryonic fibroblasts. (B-D) Confocal immunofluorescence images illustrate Kif3a (green) staining at the perinuclear region (DAPI staining for the nucleus, blue), and at the base of the primary cilium (Arl13b staining for the primary cilium, magenta) in Kif3a WT and T674A mutant mouse embryonic fibroblasts. Scale bar, 3 μm.

**Figure 3:**
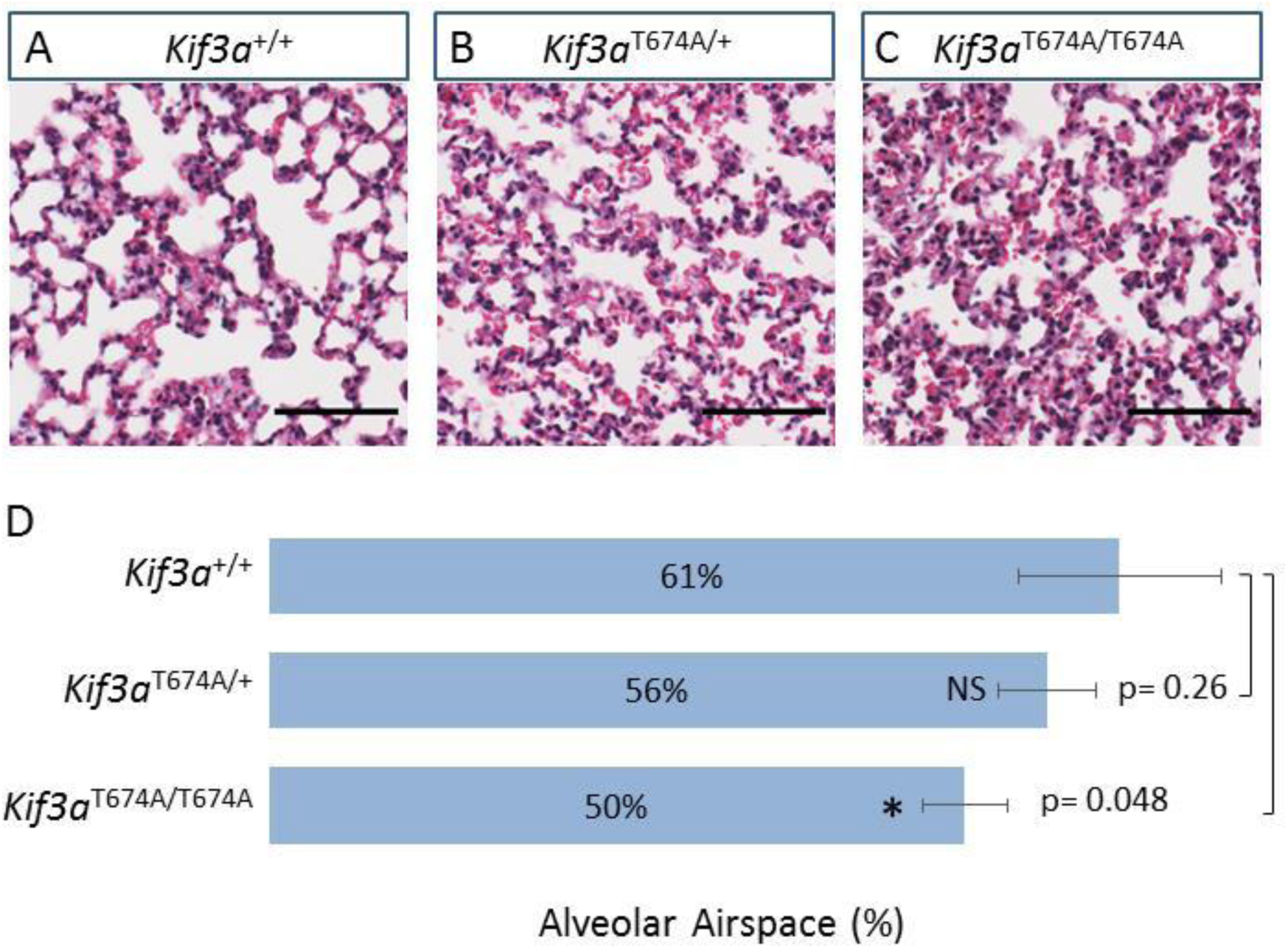
Kif3a T674A homozygous mutant lungs exhibit mildly reduced alveolar airspace. (A-C) H&E stained lung tissue sections of young adult Kif3a wild type and T674A mutant mice (n=4 per genotype, 4 weeks old). Scale bar, 80 μm. (D) Air sac area measurements for the entire lung tissue of young adult mice (n=4, 4 weeks old), excluding non-alveolar associated areas including bronchi and blood vessels. Shown are average ± SD, *P < 0.05, NS = not significant.

### 2.2 Cilia phenotype of *Kif3a* T674A knock-in cells

We investigated the impact of this phosphosite mutation in Kif3a on cilia length and formation. We isolated mouse embryonic fibroblasts (MEFs) from Kif3a WT and T674A mutant embryos. We immunostained MEFs with cilia marker Arl13B, calculated percent of ciliated cells and then measured cilia length. Our results indicate that about 60% of wild type MEF cells were ciliated under normal growth conditions, and the average cilia length was 3.7 μm (Fig. 5, 6). By comparison, MEFs bearing *Kif3a* homozygous T674A mutation showed an insignificant change in percent of ciliation but an 8% increase in average cilia length (Fig. 5, 6). MEFs bearing *Kif3a* heterozygous T674A mutation, however, were not significantly different from WT MEFs in cilia number and cilia length.

**Figure 4:**
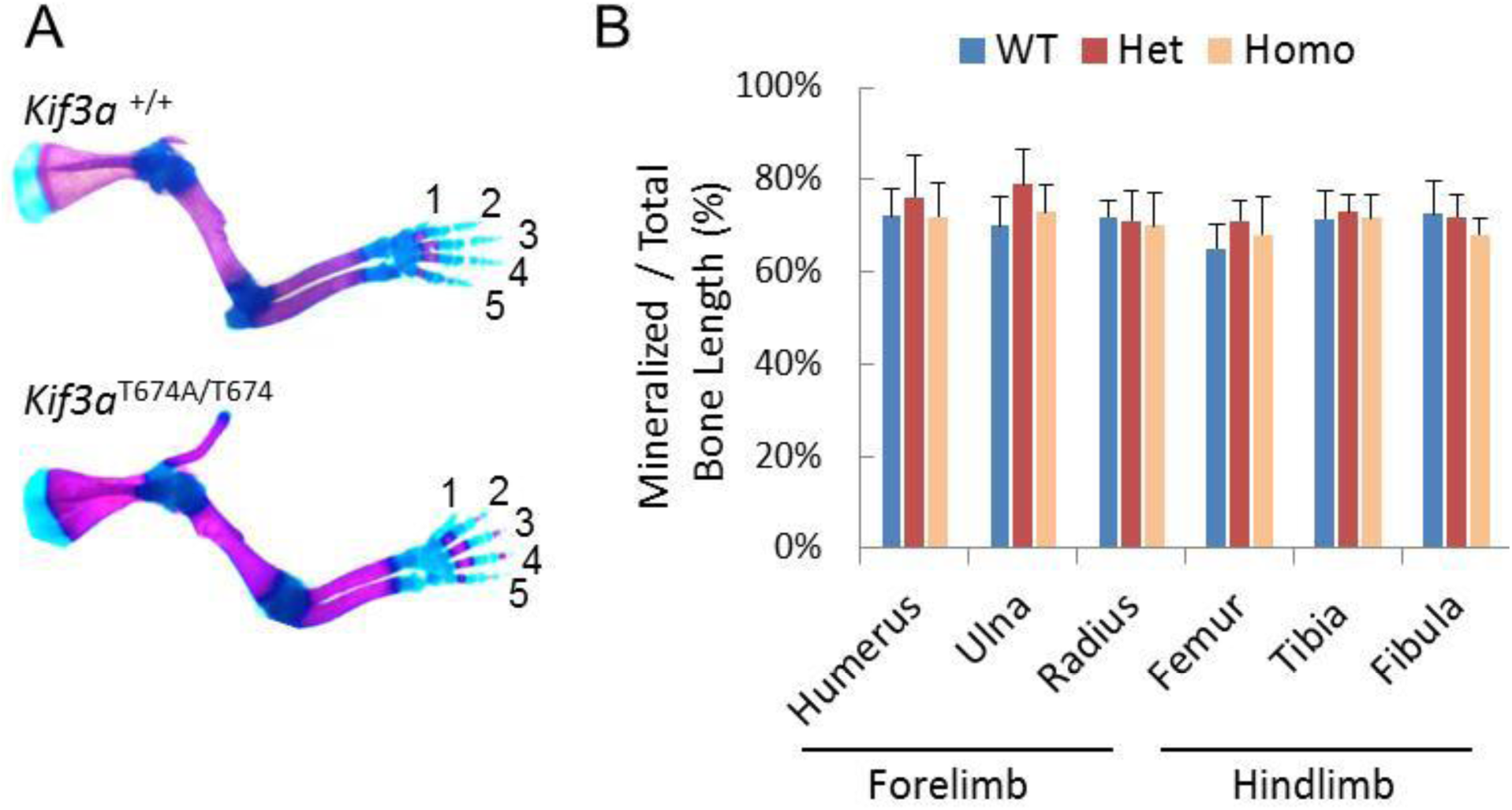
Kif3a T674A mutant limbs exhibit normal bone and digit structure. Shown in (A) are representative images of Kif3a wild type and T674A homozygote forelimb stained in Alizarin red S (calcified tissue in red) and Alcian blue (cartilage tissue in blue), and in (B) are quantification data of bone length (mineralized bone versus total bone length) of Kif3a WT and T674A mutant littermates of postnatal day 2 (average ± SD, n=4 mice per genotype).

**Figure 5:**
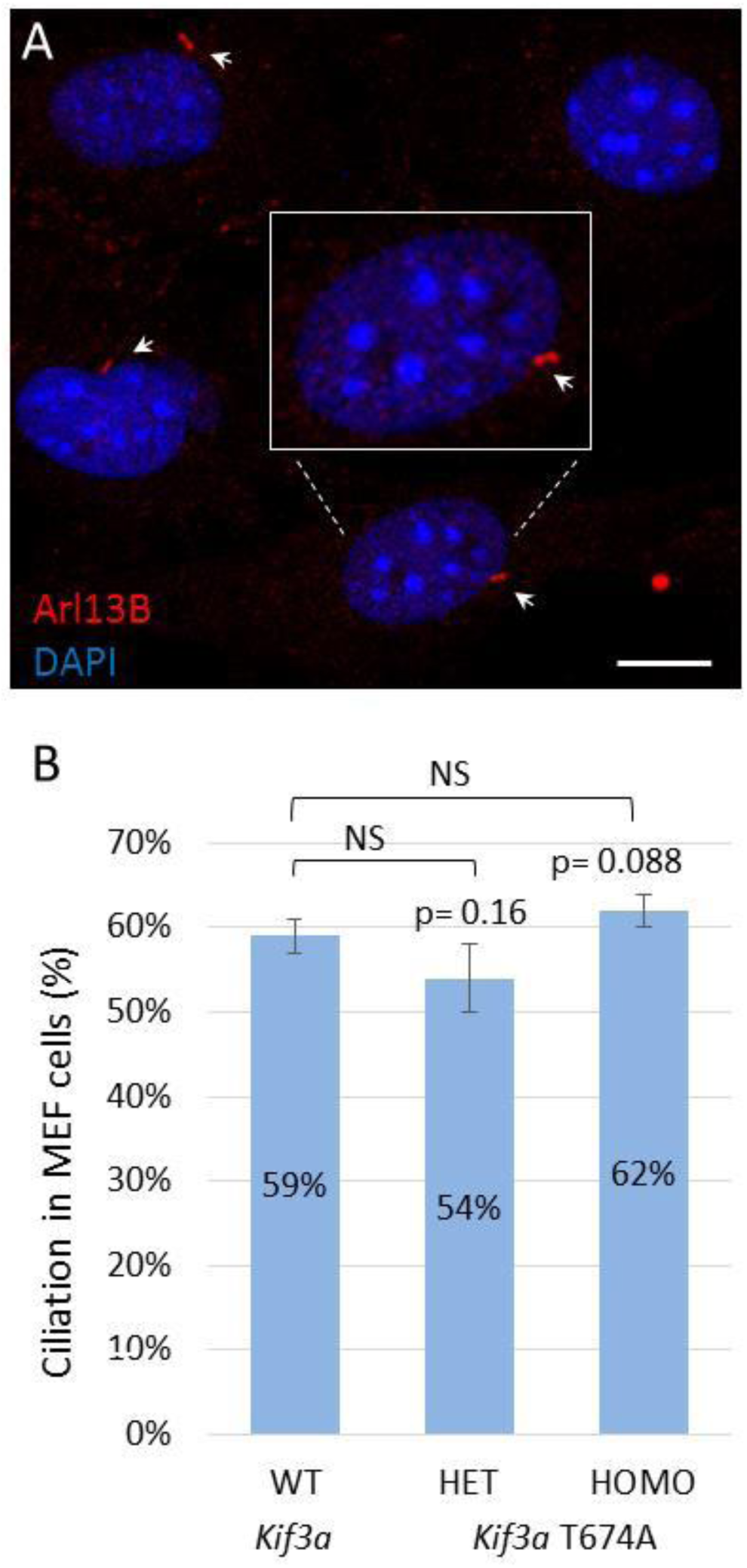
Kif3a T674A mutant cells show insignificant change in the percent of ciliation. (A) A representative confocal immunofluorescence image shows ciliated versus non-ciliated mouse embryonic fibroblasts. Primary cilia are labelled by Arl13B (red, arrow), and nucleus by DAPI (blue). Scale bar, 10 μm. (B) Measurement of percent of ciliation in Kif3a WT and T674A heterozygous (HET) and homozygous (HOMO) mutant cells. Shown are average ± SD, n=140 cells per genotype, NS = not significant.

**Figure 6:**
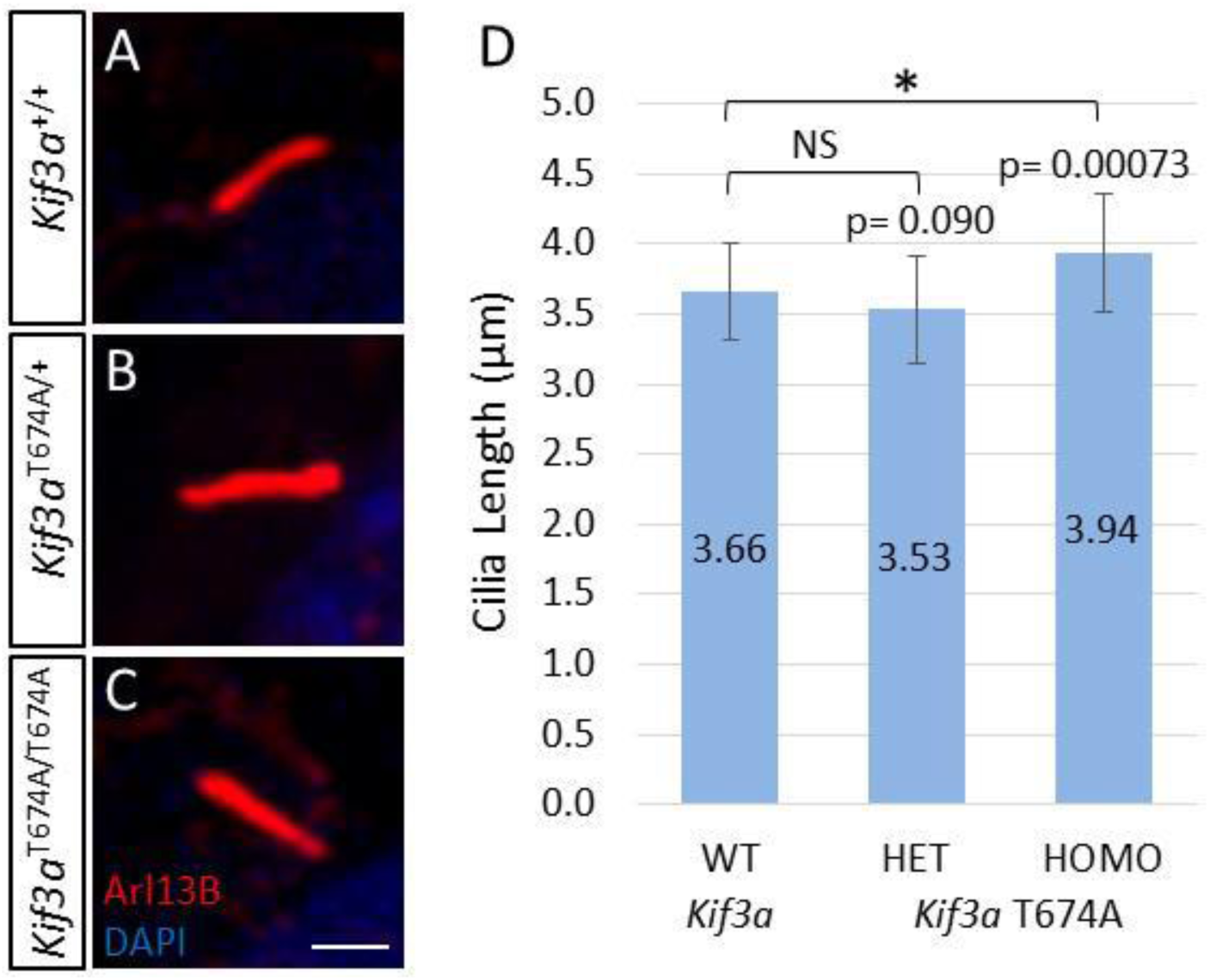
Kif3a T674A homozygous mutant cells show a moderate increase in cilia length. (A-C) Confocal immunofluorescence images show primary cilia (Arl13B, red) and nucleus (DAPI, blue) in Kif3a WT and T674A mutant mouse embryonic fibroblasts. Scale bar, 2 μm. (D) Measurement of cilia length in Kif3a WT and T674A heterozygous and homozygous mutant cells. Shown are average ± SD, n=45 cilia per genotype, *P < 0.05, NS = not significant.

We immunostained MEFs to examine distribution patterns of three ciliary proteins: intraflagellar transport protein 88 (IFT88), adenylate cyclase 3 (AC3), and GLI family zinc finger protein 2 (GLI2). IFT88 showed two basic ciliary distribution patterns: enrichment at the base and along the axoneme, and enrichment at the base and the tip (Fig. 7A). This localization pattern of IFT88 was not significantly altered by Kif3a T674A mutation (Fig. 7B). Both AC3 and GLI2 were distributed along the entire axoneme of the cilium, showing no obvious enrichment at the base or the tip. Like IFT88, the ciliary localization patterns of AC3 and GLI2 were not considerably different between Kif3a WT and T674A mutant cells (Fig. 7C). These data together indicate that although Kif3a T674A mutant cilium was slightly longer, the distribution of selected ciliary proteins was not grossly different from that in wild type cilia.

**Figure 7:**
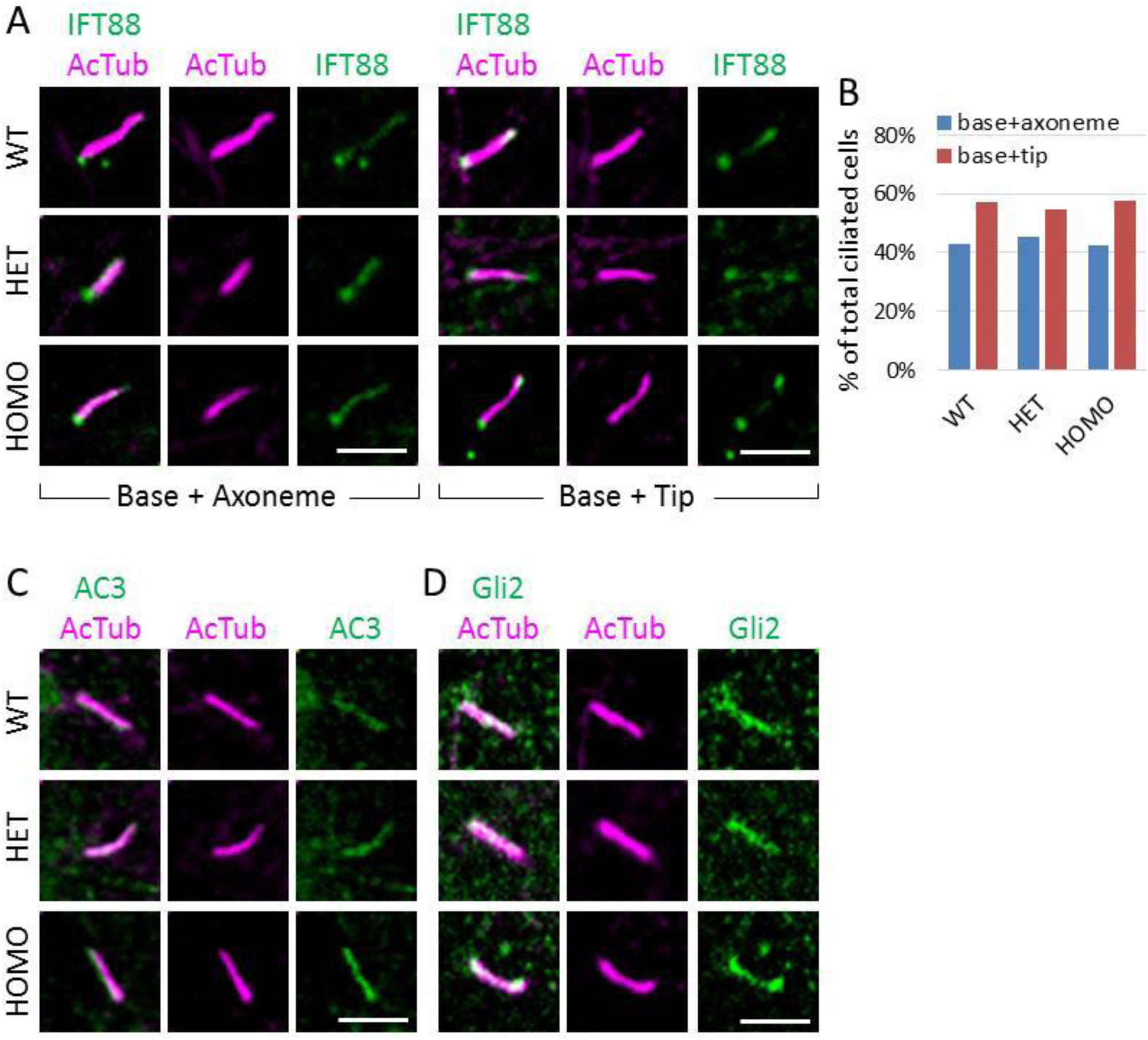
Kif3a wild type and T674A mutant cells exhibit similar distribution patterns of several ciliary proteins. Confocal immunofluorescence images illustrate ciliary localization of IFT88 (A), AC3 (C), and GLI2 (D) in Kif3a WT and T674A mutant mouse embryonic fibroblasts. Scale bar, 3 μm. The percent of two IFT88 ciliary localization patterns between different *Kif3a* genotypes is shown in (B).

## 3. DISCUSSION

By knocking-in a mutation at Thr-674 in Kif3a that prevents phosphorylation by Cilk1 at this site, our study addresses whether, and to what extent, disruption of phosphorylation of a specific substrate of CILK1 can recapitulate the profound developmental and ciliary phenotypes of ciliopathies linked to *CILK1* loss of function. Our results demonstrate that without phosphorylation of Kif3a at Thr674 there is only a mild impact on the primary cilia and tissue development. This suggests that CILK1 requires other substrates in addition to KIF3A to control ciliary structure and function and tissue development.

While inactivating mutations of *Cilk1* cause significant reduction of cilia number and change in cilia length, the *Kif3a* T674A mutation produces only moderate effects on cilia length and no change in cilia number in MEFs. This disparity in phenotypes between Cilk1 and Kif3a mutants suggests that phosphorylation of Kif3a-Thr674 plays a limited role in CILK1 regulation of cilia. Our results here underscore the importance of testing other candidate substrates of CILK1, such as Scythe and mTORC1,^22-24^ for their individual and synergistic contributions to the control of cilia assembly and maintenance.

Lung and cilia phenotypes were only detected in mice and MEFs carrying both *Kif3a* T674A mutant alleles, indicating that the phosphosite-deficient mutation T674A in *Kif3a* is recessive. *Cilk1* null or R272Q mutations are also recessive in producing ciliopathy phenotypes. These results together suggest that the functional allele of *Cilk1* or *Kif3a* is haplosufficient to contribute to the maintenance of cilia length and alveolar airspace.

Our findings add new insights into the role of KIF3A C-terminal phosphorylation in the control of cilia biogenesis and cilia length. Phosphorylation of S687/T692/S696 in KIF3A C-terminus enhances cargo-binding and transport activities, based on changes due to mutation of all three sites. In contrast, our data showing that lack of Kif3a phosphorylation at Thr674 induces longer cilia implicates an inhibitory effect of Kif3a phosphorylation on cilia growth. Do these nearby phosphosites in KIF3A C-terminus have synergistic or opposing effects on cilia? How these multiple sites coordinately regulate cilia transport, structure, and function under both physiological and pathological conditions? These questions merit further consideration and experimentation.

The precise mechanism underpinning the regulation of cilia length by KIF3A-Thr672 phosphorylation is unclear. Although our study did not reveal any gross mislocalization of several ciliary proteins, more detailed and quantitative analyses are required to provide definitive evidence showing the impact of KIF3A-Thr672 phosphorylation on ciliary transport. To follow up on our phenotypic analysis presented here, we will further investigate several possible mechanisms that may link this phosphorylation event to the regulation of cilia transport and cilia length, including any potential effects on KIF3A interactions with KIF3B and KAP in the Kinesin-2 complex, KIF3A motor activity, and KIF3A cargo-binding ability.

## 4. EXPERIMENTAL PROCEDURES

### 4.1 *Kif3a* T674A knock-in mouse strain

All procedures involving animals were performed in accordance with ethical standards in an animal protocol that was approved by the Institutional Animal Care and Use Committee. The CRISPR-assisted technology was used to generate the *Kif3a* T674A knock-in mouse. A sgRNA (Fig. 1A) is selected based on the search via the CRISPR guide design algorithm CRISPOR (http://crispor.tefor.net/). The T674A (ACC>GCA) point mutation and a silent mutation (CCA>CCT) for creating a StuI restriction cutting site near the T674A mutation were introduced into the exon 17 of the wild-type *Kif3a* allele to generate *Kif3a*/T674A repair template (ssODN). Reagents crRNA, tracrRNA, Cas9 and ssODN were purchased from IDT (Coralville, IA). Reagents crRNA and tracrRNA were diluted to 1 μg/μl in RNase-free microinjection buffer (10 mM of Tris-HCl, pH 7.4, 0.25 mM of EDTA). 5 μl crRNA and 10 μl tracrRNA were mixed and annealed in a thermocycler by heating the mixture to 95°C for 5 minutes and ramped down to 25°C at 5°C/min. Ribonucleic protein (RNP) complex was formed by mixing and incubating Cas9 at 0.5 μg/μl with crRNA/tracrRNA at 0.15 μg/μl in RNase-free microinjection buffer at 37°C for 10 minutes. Reagent ssODN containing the desired amino acid substitutions was added at 0.5 μg/μl concentration. Fertilized eggs were collected from B6SJLF1 females mated with males. RNP and ssODN were co-delivered into fertilized eggs by electroporation with BioRad Gene Pulser electroporator (six pulses at 30 V for 3 msec, separated by a 100 msec interval). The zapped zygotes were cultured overnight in KSOM medium (EMD Millipore, Billerica, MA) at 37°C in 5% CO2. The next morning, zygotes that had reached the two-cell stage were implanted into the oviducts of pseudo pregnant foster mothers of ICR strain (Envigo, Indianapolis, IN). Pups born to the foster mothers were screened using tail snip DNA by PCR genotyping and restriction digestion analysis followed by Sanger’s sequencing. Germline transmission of the desired alleles was confirmed by breeding the founders with wild type C57BL/6J mice. Animals were housed in a temperature-controlled colony room on a 12-hour light cycle, and had access to food and water ad libitum. Mice were euthanized by CO2 inhalation before tissue harvest for histological and morphometric analysis.

### 4.2 Histology and morphometric analysis

Lung tissues were isolated from young adult mice of approximately four weeks old for morphometric analysis. Lungs were extracted, fixed and embedded as previously described.^11^ Whole lung cross sections were mounted and stained with H&E (Hematoxylin and Eosin) for histological analysis. An Aperio ImageScope Slide Scanning Microscope (Leica Biosystems, Wetziar, Germany) was used to process the stained tissues and Fiji/ImageJ (version 1.52t) was used to process and analyze the alveolar air space of WT and *Kif3a* T674A mutant lungs. Each lung was separated into smaller fields wherein the images were prepared for analysis by removing field areas of non-lung regions, blood vessels, and bronchioles so that only the alveolar-associated air space and lung tissue were quantified. These were then converted into binary images in ImageJ so that area of the tissue and air space could be separately quantified.

Limb bones were dissected from mice of postnatal day 2 for morphometric analysis. Bones were rinsed with PBS and then fixed in 95% ethanol for 2 days, followed by acetone overnight at room temperature. Limbs were stained with 0.03% Alcian blue solution, washed with two changes of 70% ethanol, and incubated with 95% ethanol overnight. Limbs were then incubated with 1% KOH for 1 h to pre-clear tissues and stained with 0.005% Alizarin red solution for 4 h. Cartilages were stained with Alcian Blue and bones were stained with Alizarin Red. After clearing, samples were placed in 100% glycerol for long-term storage and imaging (Olympus SZX12, Olympus).

### 4.3 Cell culture and Western blot

Mouse embryonic fibroblast (MEF) cells were isolated from *Kif3a* wild type and T674A mutant embryos (E14.5-E15.5) and maintained at 37°C and 5% CO2 in Dulbecco’s modified Eagle’s medium supplemented with 10% fetal bovine serum, non-essential amino acids, and penicillin-streptomycin using a standard protocol.^25^

MEF cells were lysed in lysis buffer (50 mM Tris-HCl, pH 7.4, 150 mM NaCl, 1% NP-40, 2 mM EGTA, and complete protease and phosphatase inhibitors from Roche, Basel, Switzerland). Cell lysate was cleared by centrifugation. Protein extracts were boiled for 5 min in an equal volume of 2X Laemmli sample buffer (120 mM Tris-HCl, pH 6.8, 4% SDS, 20% glycerol, 10% β-mercaptoethanol, 0.02% bromophenol blue) and loaded on an SDS-gel. Samples were transferred to a PVDF membrane and blocked for one hour in 5% dry milk before incubation with KIF3A (D7G3) rabbit monoclonal antibody (Cell Signaling Technology, Danvers, MA, #8507) in TBS containing 0.1% Tween-20 and 5% bovine serum albumin (BSA) overnight at 4°C. This was followed by extensive rinses and one-hour incubation with horseradish peroxidase (HRP)-conjugated secondary antibody. Chemiluminescent signals were developed using Millipore Immobilon ECL reagents (EMD Millipore).

### 4.4 Confocal immunofluorescence microcopy and cilia analysis

Mouse embryonic fibroblast (MEF) cells were grown on gelatin-coated coverslips were fixed by 4% paraformaldehyde (PFA) in PBS, rinsed in PBS, and then permeabilized by 0.2% Triton X-100 in PBS. After one hour in blocking buffer (3% goat serum, 0.2% Triton X-100 in PBS), cover slips were incubated in primary antibody at 4°C overnight followed by rinses in PBS and one hour incubation with goat anti-mouse IgG (Alexa Fluor 647-conjugated) secondary antibody (Abcam, Cambridge, MA, ab150115) and/or goat anti-rabbit IgG (Alexa Fluor 594-conjugated) secondary antibody (Abcam, Cambridge, MA, ab150084). MEF cells on cover slips were incubated with cilia marker Arl13B rabbit antibody (ProteinTech, Rosemont, IL, 17711-1-AP) for analysis of cilia length and prevalence. For localization analysis of ciliary proteins, additional cover slips were co-incubated with acetylated tubulin (Lys40) mouse monoclonal antibody (ProteinTech, Rosemont, IL, 66200-1) as well as with KIF3A (D7G3) rabbit monoclonal (Cell Signaling Technology, Danvers, MA, #8507), IFT88 rabbit polyclonal (ProteinTech, Rosemont, IL, 13967-1-AP), ADCY3 rabbit polyclonal (ProteinTech, Rosemont, IL, 19492-1-AP), or Gli2 rabbit polyclonal antibody (GeneTex, Irvine, CA, GTX46056). After extensive rinses, slides were mounted in antifade reagent containing DAPI (4’, 6-diamidino-2-phenylindole) for imaging via ZEISS LSM 700 Confocal Microscope.

Zen 2009 program was used with the confocal Laser Scanning Microscope 700 (LSM 700) to collect z stacks at 0.5 μm intervals to incorporate the full axoneme based on cilia marker Arl13b staining. All cilia were then measured in Fiji/ImageJ (version 1.52t) via a standardized method based on the Pythagorean Theorem in which cilia length was based on the equation L2 = z2 + c2, in which “c” is the longest flat length measured of the z slices and “z” is the number of z slices in which the measured cilia was present multiplied by the z stack interval (0.5 μm). Z stacks were used to measure cilia lengths as described, the number of ciliated cells, as well as for qualitative analysis of localization patterns of Ift88, Adcy3, Gli2, and Kif3a.

### 4.5. Statistical analysis

Quantified experimental data between different genotypes were analyzed and compared by an unpaired, two tailed Student *t*-test. Data were reported as average ± standard deviation (SD). P-values less than 0.05 were considered as significant.

## ACKNOWLEDGEMENTS

Z.F. and D.L.B. conceived the project. W.X., Z.F., E.J.W. designed, constructed, and evaluated the mouse model. C.D.G., E.J.W., L.J., S.A., Z.F., D.L.B., X.L. performed experiments and conducted data curation and analysis. Z.F., D.L.B., L.J., and X.L. contributed essential reagents/tools and acquired funding support. Z.F., C.D.G., W.X., and L.J. contributed to the original writing, D.L.B. contributed to the editing of the manuscript.

We thank our colleagues at UVA Genetically Engineered Murine Model Core, Research Histology Core, and Advanced Microscopy Facility for technical support. This work was supported by NIGMS grant GM127690 to Z.F., NIAMS grant AR064792 to X.L., NCI CCSG P30CA044579 to UVA School of Medicine research cores, University of Virginia Harrison undergraduate research grant to E.J.W., and the F. Palmer Weber endowed professorship to D.L.B.

## FOOTNOTES

CILK1, ciliogenesis associated kinase 1; ICK, intestinal cell kinase; IFT, intraflagellar transport; KIF3A/3B, kinesin family member 3A/3B; KAP, kinesin associated protein; IFT88, intraflagellar transport 88; ADCY3, adenylate cyclase 3; GLI2, GLI family zinc finger protein 2; MEF, mouse embryonic fibroblast

